# Genetic diversity assessment of Pacific oyster *Magallana gigas* (*Crassostrea gigas*) populations from the two southern coastal farms in South Korea

**DOI:** 10.1101/2022.05.30.493984

**Authors:** Biet Thanh Tran, Su-Jin Park, Hong Keun Park, Dongjin Park, Youn Hee Choi

## Abstract

South Korea is among the major producers of the Pacific oyster, *Magallana gigas* (*Crassostrea gigas*), which is one of the most valued aquaculture species. Since the early 1990s, climatic and anthropogenic factors have incurred the reduction of their wild seeds, whereby the dependence on hatchery-produced seeds has constantly increased in South Korea, thus raising concerns about losing genetic diversity and accelerating genetic deterioration. To better understand their genetic make-up, we assessed the genetic diversity of *M*. *gigas* populations from two farms (Tongyeong and Gadeokdo) in the southern coast, where about 80% of the cultivated oysters in Korea are produced. Tongyeong showed slightly higher diversity than Gadeokdo, but both populations had a similar genetic structure characterized by low nucleotide diversity. Comparative haplotype analyses provided data supporting unique genetic features of the populations that include (1) weak genotype-locality relationship, (2) low levels of gene flow between populations, and (3) seasonal fluctuation of genetic variation within a population. Furthermore, the highly alike haplotype network patterns were observed between the wild and farm populations as well as among the populations in neighboring countries, which suggests that the genetic structure is conserved between wild and hatchery populations, and geographic proximity has minimal influence on the genetic composition. These results warrant further study in biological and ecological contexts and will be invaluable in formulating genetic monitoring and sustainable long-term management of *M. gigas*.

## Introduction

The Pacific oyster, *Magallana gigas*, formerly *Crassostrea gigas*, is a seasonal delicacy beloved all over the world and has become one of the globally important aquaculture species [1]. It is also reputable for its high nutritional quality and health benefits, therefore it has been dubbed “Milk of Sea” [2, 3]. In Korea, the yearly production of *M. gigas* exceeds 300,000 metric tons, making the country the second largest producer in the world as of 2019 [4]. The Korean oyster farming used to heavily rely on the collection of natural spats that accounted for 90% for the domestic oyster seed demand, while the remaining 10% was hatchery-produced seeds [5]. However, changing climates and augmenting anthropogenic activities have significantly reduced natural *M. gigas* seedlings and led to the point where seeds fell short since 1992 [6]. In response, the farms turned their eyes to hatchery-produced seeds whose weight in the seed market has subsequently increased [7]. New hatchery-propagated stocks could be isolated from the wild populations for genetic improvement, and yet seeds harvested at the limited number of hatcheries would eventually put the *M. gigas* populations in Korea at risk of inbreeding and genetic deterioration [8]. Over the course of the successive selection process, the seed stocks are likely to lose genetic diversity as a result of founder effects and random genetic drift. Therefore, the oysters might not carry fitness-related characteristics they need after being transferred to grow-out areas [9]. This process has been well-documented in many aquaculture organisms where they had lowered genetic diversity compared with their wild progenitors [10, 11, 12].

In 2007, the 10,900 tons of crude oil released from the tanker, MV Hebei Spirit, covered 375km of shoreline of the western coast of Korea [13]. The toxicity of the oil wiped out most oysters at more than 90% of the oyster farms in that region, triggering the displacement of the natural spats by the ones from southern areas [14]. In consequence, this accident (in addition to the climatic and anthropogenic interference) further depleted the wild stock. Thereafter, a large amount of seed was imported from Japan and the United States in an effort to replenish the drastic shortage of seeds [6]. The above chronologically deteriorating circumstances raised necessity to look into the genetic composition of *M. gigas* populations in South Korea, but there is little information on the genetic variation and structure, particularly in populations at hatchery farms.

Mitochondrial DNA (mtDNA) analysis is an effective approach to examine divergences in animals by virtue of its small size, maternal and non-recombining mode of inheritance, rapid evolution, extensive intraspecific polymorphisms, etc [15, 16]. Among mtDNA markers, cytochrome oxidase *c* subunit I (mtCOI) is one of the most conserved protein-encoding genes in the mitochondrial genome [17]. The effectiveness of mtCOI analysis to study genetic diversity and genetic structure of marine mollusk populations has been charted in *Tegillarca granosa*, *Potamocorbula laevis*, *Meretrix petechialis*, etc [18, 19, 20]. Li et al. [21] interrogated mtCOI to analyze the population structure and demographic history of *M. gigas* in the northwestern Pacific region. In Korea, although a few studies have examined the genetic diversity of *M. gigas* using molecular-based methods [22, 23], it is still insufficient, given the economic value of this species.

The present study investigated the mtCOI gene to elucidate the genetic variation and population structure of *M. gigas* from two aquaculture farms in Tongyeong and Gadeokdo in the southern coast, where about 80% of the cultivated oysters in Korea are produced [24, 25], with the aim of providing fundamental information to sustainably maintain the cultivated stock.

## Materials and Methods

### Sample collection

Forty oysters each were obtained from aquaculture farms in Tongyeong (TY, 34°52’38.4”N 128°14’40.8”E) and Gadeokdo (GD, 35°04’19.0”N 128°50’28.6”E) with ten collected at each site in October 2020 and January, April, and July 2021 (Fig 1). They were immediately transferred on ice to the Laboratory of Physiology, Pukyong National University, Busan, Korea. Their fresh mantle edges were dissected and stored in 100% ethanol at room temperature until DNA isolation.

**Fig 1.**
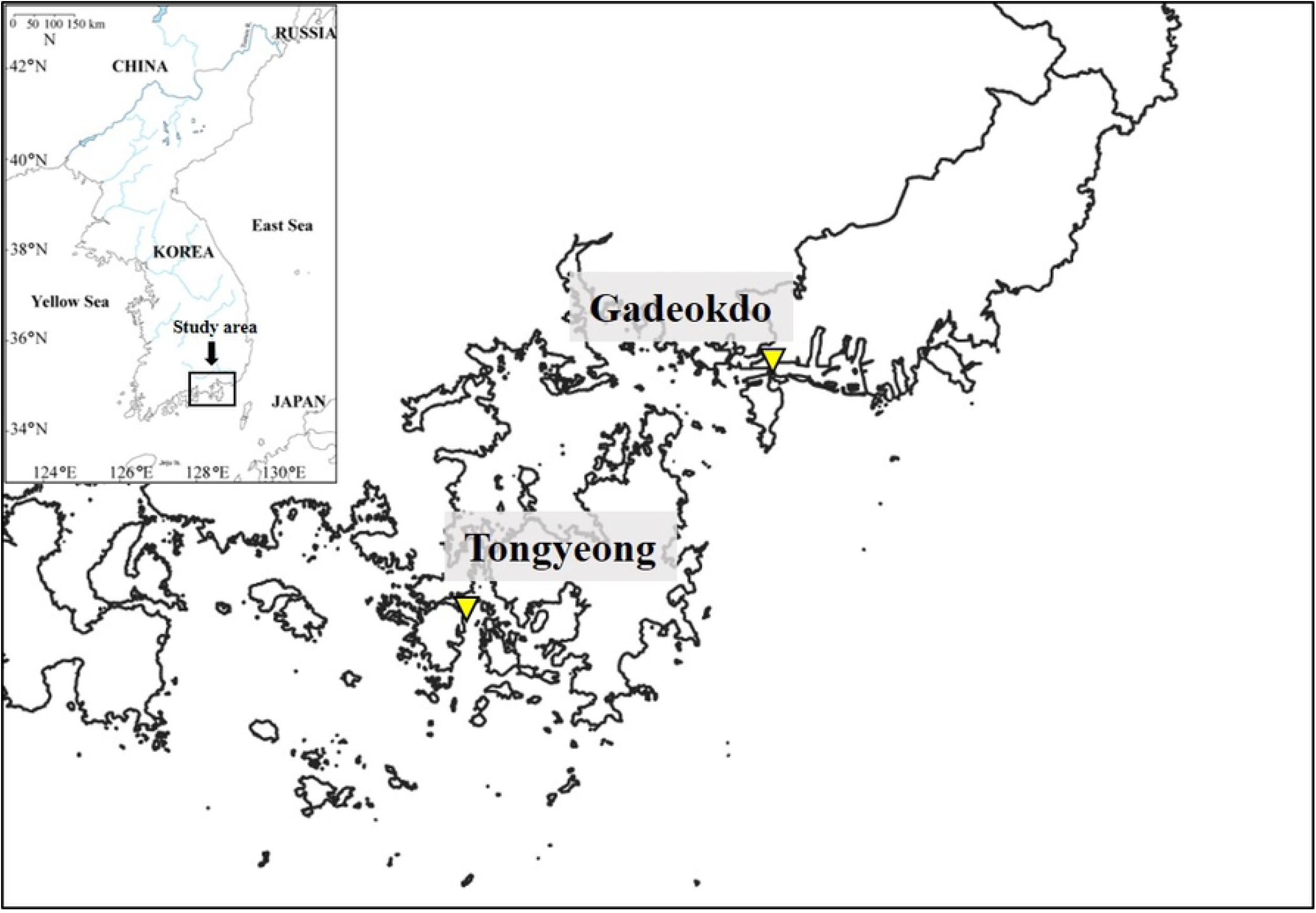
Pacific oyster, *M. gigas*, sampling sites in Tongyeong (TY) and Gadeokdo (GD), South Korea.

### DNA extraction

The excised samples were incubated with 600 μL of TNES-urea buffer (10 mM Tris-HCl, pH 8.0; 125 mM NaCl; 10 mM EDTA, pH 8.0; 0.5% SDS; 6 M urea) and 10 μL proteinase K (20 μg/mL) at 60°C for at least 12 h, and total genomic DNA (gDNA) was extracted using the phenol/chloroform method [26]. The extracted gDNA was dried at room temperature, resuspended in TE buffer (10mM Tris-HCl; 1 mM EDTA, pH 8.0), and stored at −20°C for later use. Its quantity and quality were checked using a spectrophotometer (DS-11; DeNovix, DE, Wilmington, USA).

### Primer preparation

To design a pair of primers for polymerase chain reaction (PCR), we retrieved mtCOI sequences from GenBank (https://www.ncbi.nlm.nih.gov/) and aligned them using ClustalW in BioEdit 7.2 [27]. After comparing the aligned nucleotide matrix, conserved regions of the *Magallana* sequences were selected as forward and reverse primers. Their melting temperatures and secondary structures were checked with the web service Sequence Manipulation Suite [28] before oligonucleotide synthesis.

### PCR amplification and sequencing

PCR was conducted in a 20-μL reaction using *AccuPower*^®^ PCR PreMix (Bioneer, Daejeon, South Korea) with 0.2 μM of the forward [CRA-MT-18171f (5’-TTTCGCACTGGGCAGGTAT-3’)] and reverse [CRA-MT-1865r (5’-ATTTCGCACTGGGCAGGTAT-3’)] primers and 1 μL of gDNA template (20 ng/μL). The DNA was amplified using the ProFlex PCR System (Thermo Fisher Scientific, Waltham, MA, USA) at 95°C for 5 min, followed by 35 cycles at 95°C for 30 s, 55°C for 30 s, and 72°C for 2 min, with a 5-min final extension at 72°C. The PCR product was electrophoresed in 1.5% agarose gel stained with ChamelGreen^™^ I (SFC Probes, South Korea) followed by gel purification with an *AccuPrep*^®^ PCR Purification Kit (Bioneer, Daejeon, South Korea). The purified DNA was subsequently sequenced at Macrogen (Seoul, South Korea).

### Data analysis

The raw sequence data were imported to Sequencher v. 5.4.5 (Gene Codes, Ann Arbor, MI, USA) to check the base calls on the chromatograms manually. We aligned all sequences using the Clustal W function in BioEdit 7.2.5 with the default parameters. After manual correction of obvious misalignments, a concatenated nucleotide sequence matrix of 863-bp mtCOI fragments from the 80 *M. gigas* specimens from TY and GD was generated. DnaSP v. 6 [29] was employed to estimate the total number of haplotypes (H) and haplotype distribution among populations. The haplotype diversity (h), nucleotide diversity (π), and mean number of pairwise differences (k) were calculated using ARLEQUIN v. 3.5 [30].

A parsimony network was generated and plotted with TCS analysis [31] using PopART v. 1.7 [30] to evaluate the genealogical relationship of all *M. gigas* haplotypes. The network was constructed from 80 sequences from the TY and GD populations and 117 sequences (retrieved from GenBank) from China, Japan, Korea, Canada, Taiwan, Brazil, Mexico, Netherlands, and the United States (Table 1).

**Table 1.**
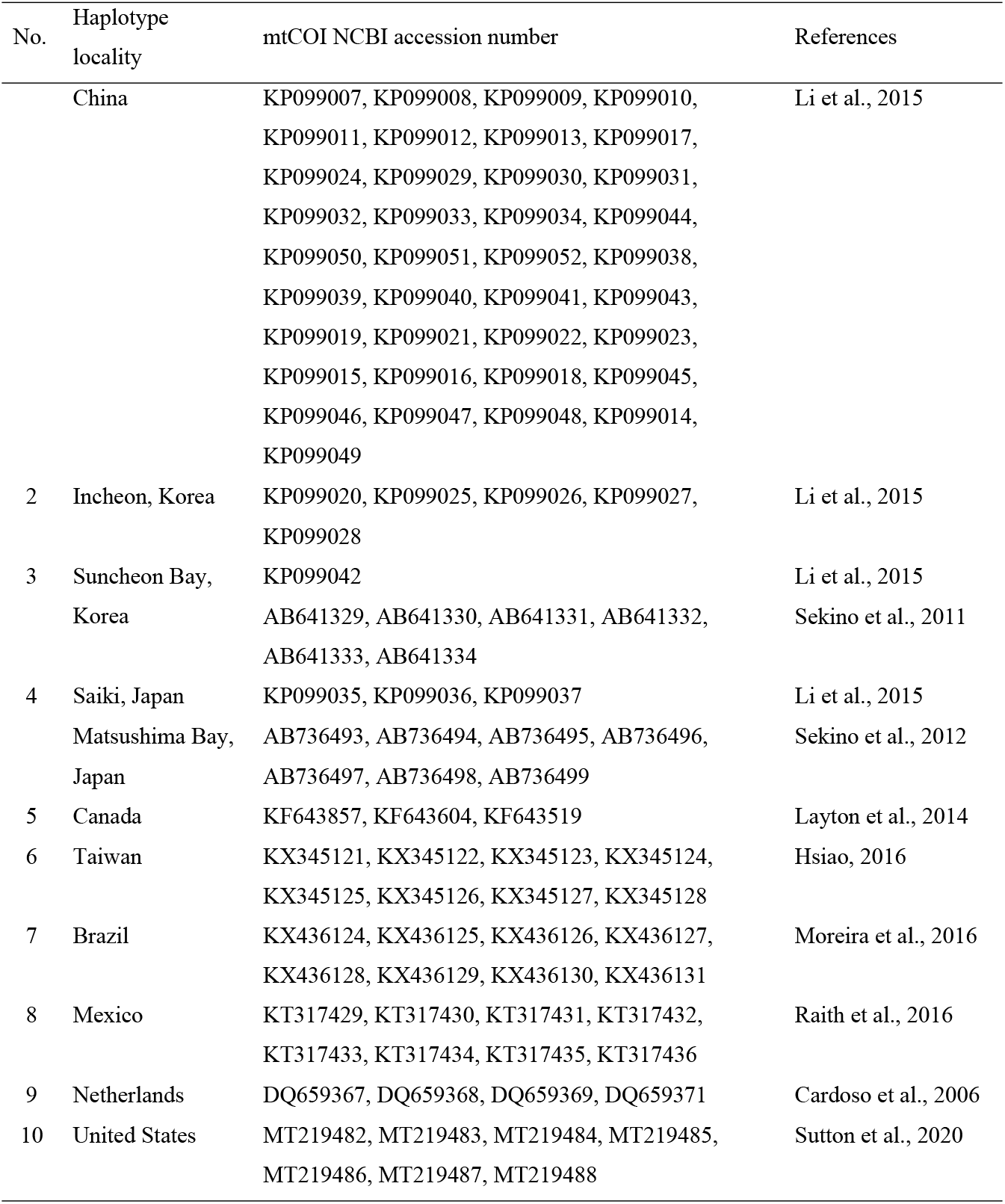
mtCOI sequences of various *M. gigas* populations retrieved from GenBank.

| No. | Haplotype locality | mtCOI NCBI accession number | References |
| --- | --- | --- | --- |
|  | China | KP099007, KP099008, KP099009, KP099010, KP099011, KP099012, KP099013, KP099017, KP099024, KP099029, KP099030, KP099031, KP099032, KP099033, KP099034, KP099044, KP099050, KP099051, KP099052, KP099038, KP099039, KP099040, KP099041, KP099043, KP099019, KP099021, KP099022, KP099023, KP099015, KP099016, KP099018, KP099045, KP099046, KP099047, KP099048, KP099014, KP099049 | Li et al., 2015 |
| 2 | Incheon, Korea | KP099020, KP099025, KP099026, KP099027, KP099028 | Li et al., 2015 |
| 3 | Suncheon Bay, Korea | KP099042<br>AB641329, AB641330, AB641331, AB641332, AB641333, AB641334 | Li et al., 2015<br>Sekino et al., 2011 |
| 4 | Saiki, Japan<br>Matsushima Bay, Japan | KP099035, KP099036, KP099037<br>AB736493, AB736494, AB736495, AB736496, AB736497, AB736498, AB736499 | Li et al., 2015<br>Sekino et al., 2012 |
| 5 | Canada | KF643857, KF643604, KF643519 | Layton et al., 2014 |
| 6 | Taiwan | KX345121, KX345122, KX345123, KX345124, KX345125, KX345126, KX345127, KX345128 | Hsiao, 2016 |
| 7 | Brazil | KX436124, KX436125, KX436126, KX436127, KX436128, KX436129, KX436130, KX436131 | Moreira et al., 2016 |
| 8 | Mexico | KT317429, KT317430, KT317431, KT317432,<br>KT317433, KT317434, KT317435, KT317436 | Raith et al., 2016 |
| 9 | Netherlands | DQ659367, DQ659368, DQ659369, DQ659371 | Cardoso et al., 2006 |
| 10 | United States | MT219482, MT219483, MT219484, MT219485,<br>MT219486, MT219487, MT219488 | Sutton et al., 2020 |

The genetic structure of the *M. gigas* populations from the TY and GD was assessed by analysis of molecular variance (AMOVA) on ARLEQUIN v. 3.5. The fixation index (FST) value ranges from 0 (no differentiation between subpopulations) to 1 (complete differentiation of populations) [33]. The significance of each pairwise comparison was tested with 10,000 random replicates.

## Results

### Amplification of mtCOI gene from a chronological set of *M. gigas*

Eighty *M. gigas* samples collected from aquaculture farms in Tongyeong (TY) and Gadeokdo (GD) over the 4 consecutive seasons (i.e., 10 TY and 10 GD samples per season) were used for this study. Initially, the mtCOI gene of each individual was amplified by PCR. All samples generated a single 2000-bp amplicon, indicating the presence of an intact mtCOI gene in each individual (Fig 2). Subsequent sequencing confirmed that all of the amplicons are mtCOI-specific (data not shown).

**Fig 2.**
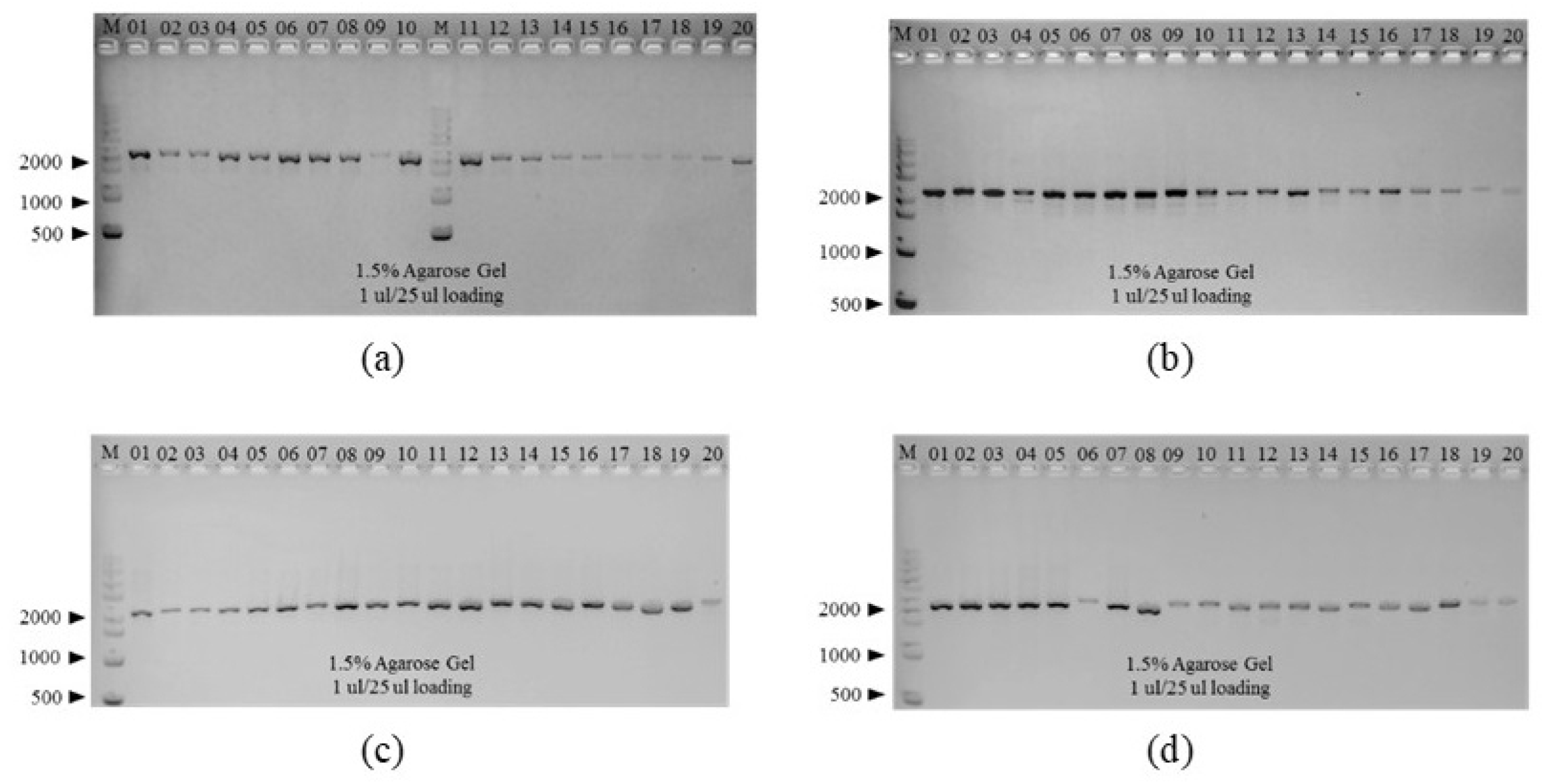
PCR amplification of mitochondrial *cytochrome c oxidase subunit 1* from *M. gigas* samples collected from TY (01 to 10) and GD (11 to 20) in the (a) fall, (b) winter, (c) spring, and (d) summer. A pair of primers, CRA-MT-18171f and CRA-MT-1865r, were used to amplify a 2000-bp fragment that includes the gene. M, 1 kb size marker.

### Genetic diversity

To characterize the genetic diversity of *M. gigas* populations from TY and GD, multiple sequence analysis was performed using a 863-bp concatenated nucleotide sequence of the mtCOI from the 80 samples. This analysis identified 27 different single nucleotide polymorphism (SNP) sites in total, consisting of 21 transitions (a/g, c/t) and 4 transversions (t/g), with no indels being detected. The translated amino acid sequences of the mtCOI fragments from the 80 *M. gigas* were found to be 100% identical, indicating that the sequence variation of *M. gigas* mtCOI is predominantly due to synonymous SNPs (data not shown).

Based on the SNP profiles, a total of 21 haplotypes were determined; 13 in TY and 9 in GD (Table 2), which indicates that the genetic diversity is slightly greater in TY (h=0.5154) than GD (h=0.4397) during the study period (Table 2)”. Haplotype 1 (HAP_1) was the most abundant haplotype shared by 58 samples (28 in TY and 30 in GD) and accounted for 72.5% of the total sample size. Other 20 haplotypes differed from HAP_1 by 1-3 SNPs. As expected, both TY and GD populations displayed low nucleotide diversity (0.000927 and 0.000689, respectively), with TY being slightly more diverse (Table 2). Consistently, the haplotype diversity of TY was higher than that of GD. The mean number of pairwise differences (k) measuring the genetic diversity within a population was higher in TY than in GD (Table 2). Overall, TY and GD populations shared a similar genetic profile characterized by low genetic diversity with the predominant HAP_1 and diverse minor haplotypes.

**Table 2.**
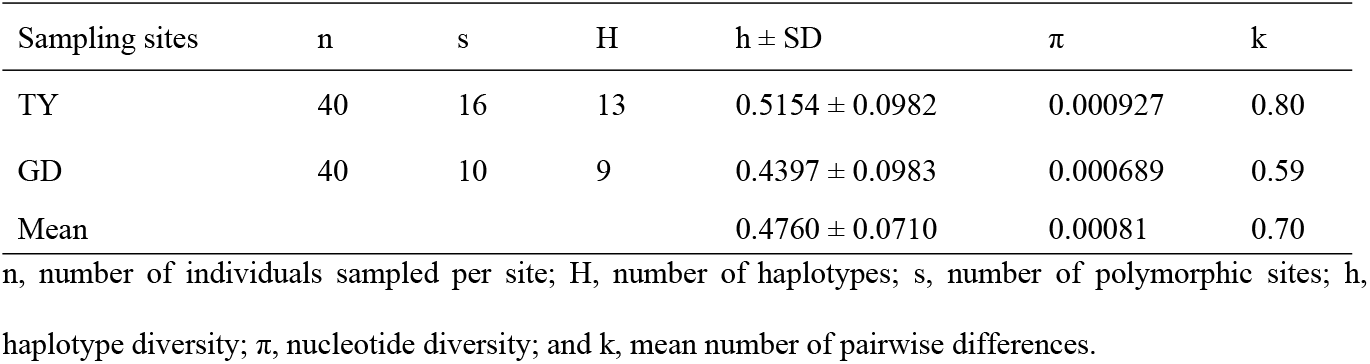
Diversity indices of *M. gigas* from TY and GD.

### Population genetic structure

AMOVA analysis was conducted to characterize the genetic structure of the 2 *M. gigas* populations. As shown in Table 3, a majority of the total molecular variance was present within populations (99.81%) rather than between the 2 populations. Accordingly, the fixation index (FST), a measure of population differentiation due to genetic structure, was 0.00188 but was not statistically significant (*p*-value = 0.49853).

**Table 3.**
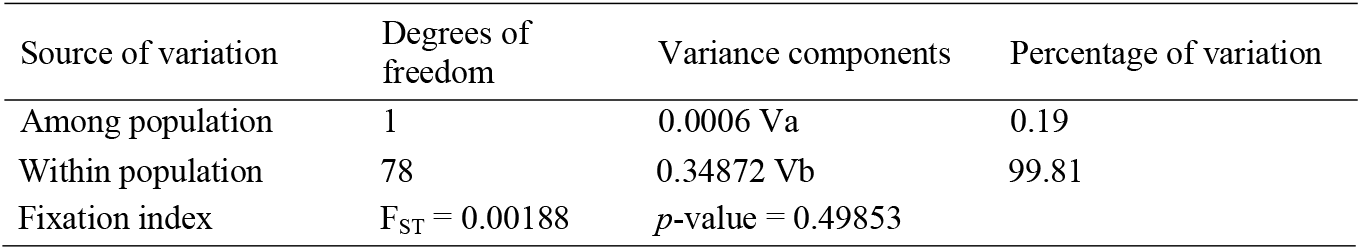
Analysis of molecular variance (AMOVA) among *M. gigas* mtCOI sequences

| Source of variation | Degrees of freedom | Variance components | Percentage of variation |
| --- | --- | --- | --- |
| Among population | 1 | 0.0006 $V_a$ | 0.19 |
| Within population | 78 | 0.34872 Vb | 99.81 |
| Fixation index | $F_{ST} = 0.00188$ | $p\text{-value} = 0.49853$ | |

### Characterization of haplotypes

To further delineate the genetic relationship of the *M. gigas populations*, a haplotype network was constructed using the 863-bp mtCOI alignment matrix. As shown in Figure 3, HAP_1 formed a core of the network with other 20 satellite haplotypes. Except for HAP_19 and HAP_20, all satellite haplotypes were shown to be a singleton with a distinct SNP profile. All satellite haplotypes differ from HAP_1 by 1-3 SNPs, resulting in the characteristic ‘star-like’ network pattern. Overall, geographical clustering between TY and GD was not observed in the network.

**Fig 3.**
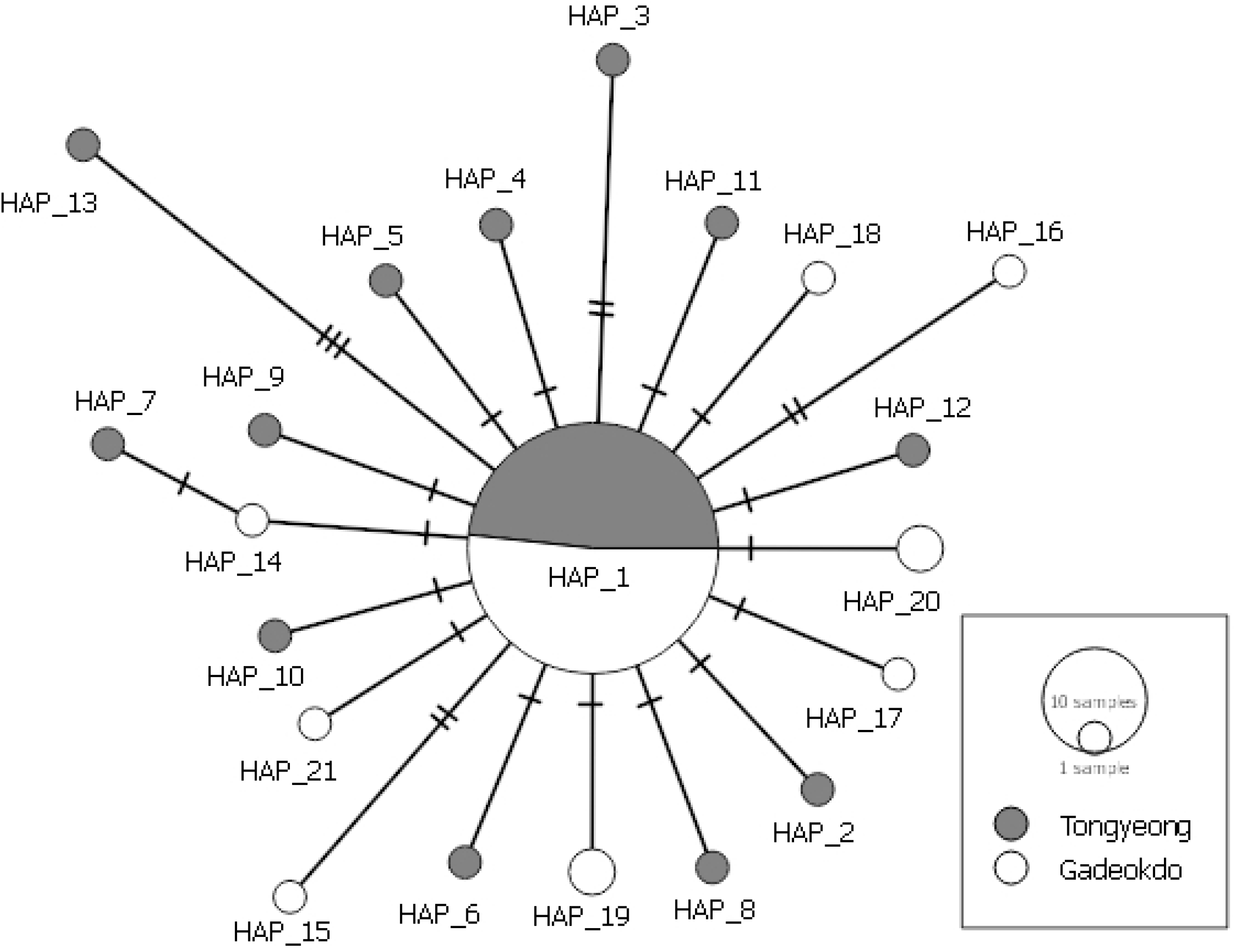
Haplotype network of mtCO1 of *M. gigas* populations from TY and GD. Each circle corresponds to a unique haplotype sized in proportion to its frequency. The hash marks on the lines indicate the number of substitutions that separate haplotypes.

In addition, the seasonal pattern of haplotypes was accessed in TY and GD populations (Table 4). As inferred from the haplotype network, 50 to 90% of sequences in each season were found to be HAP_1. Interestingly, different satellite HAPs were detected in each season, except Hap_19 and 20 which are commonly found in Spring and Summer at GD, indicating that the genetic stability is maintained by HAP_1 while frequent replacements of satellite haplotypes mainly contribute to genetic diversification in both *M. gigas* populations over time.

**Table 4.**
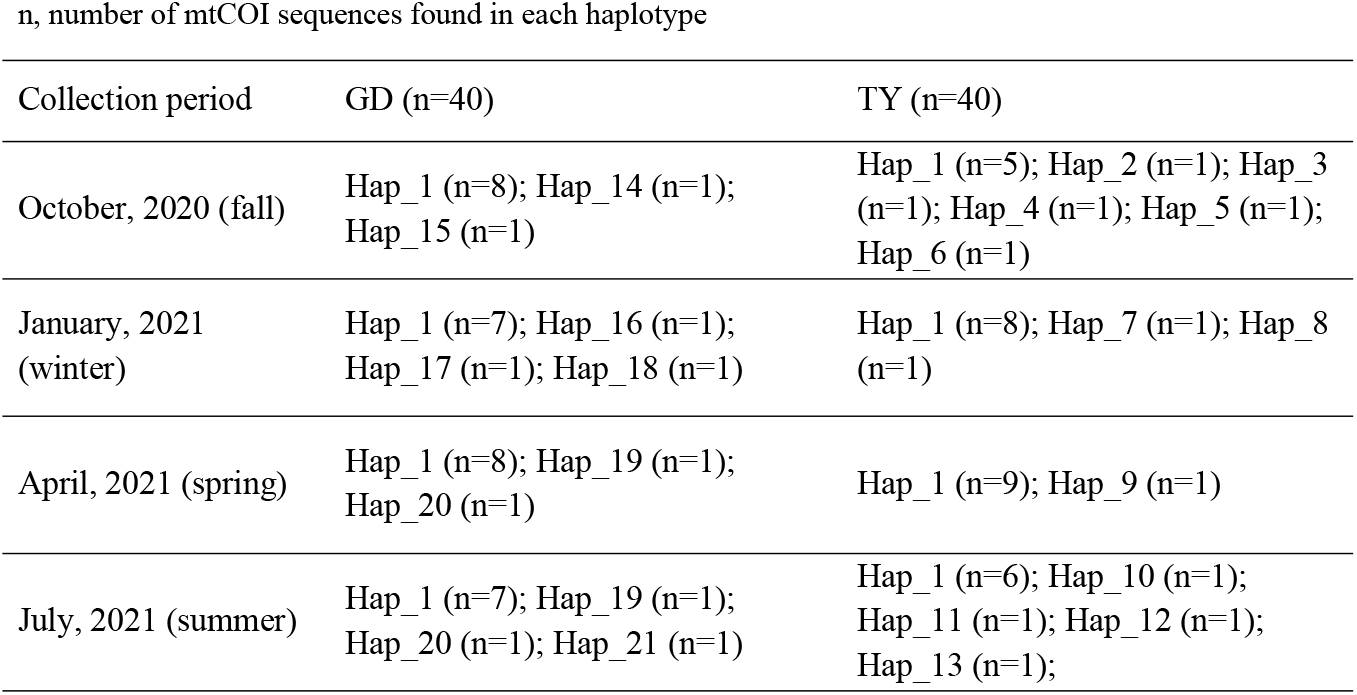
Temporal variation of haplotypes

Next, 117 mtCOI sequences of 8 different county origins and two Korean origins (Incheon and Sucheon) were retrieved from the NCBI GenBank and included in the haplotype network analysis (Table 1). Li et al. [21] previously characterized 46 HAPs in the *M. gigas* population with 12 geographical origins in the northwestern Pacific. Selected mtCOI sequences representing 37 haplotypes, including HAP_1 (KP099007), out of the 46 HAPs were included in this analysis (see Table 1). Similar to Fig 3, the haploid network consisted of the dominant HAP_1 and the satellite HAPs. While a majority of satellite haplotypes was a singleton, 11 satellite haplotypes shared by different geographical origins were identified (Fig 4). Despite the geographical proximity, only 3 haplotypes (i.e., HAP_3, HAP_11, and HAP_14) were shared by Chinese and Korean *M. gigas* populations. No significant relationship was observed between the Korean origins (TY, GD, Incheon, and Sucheon).

**Fig 4.**
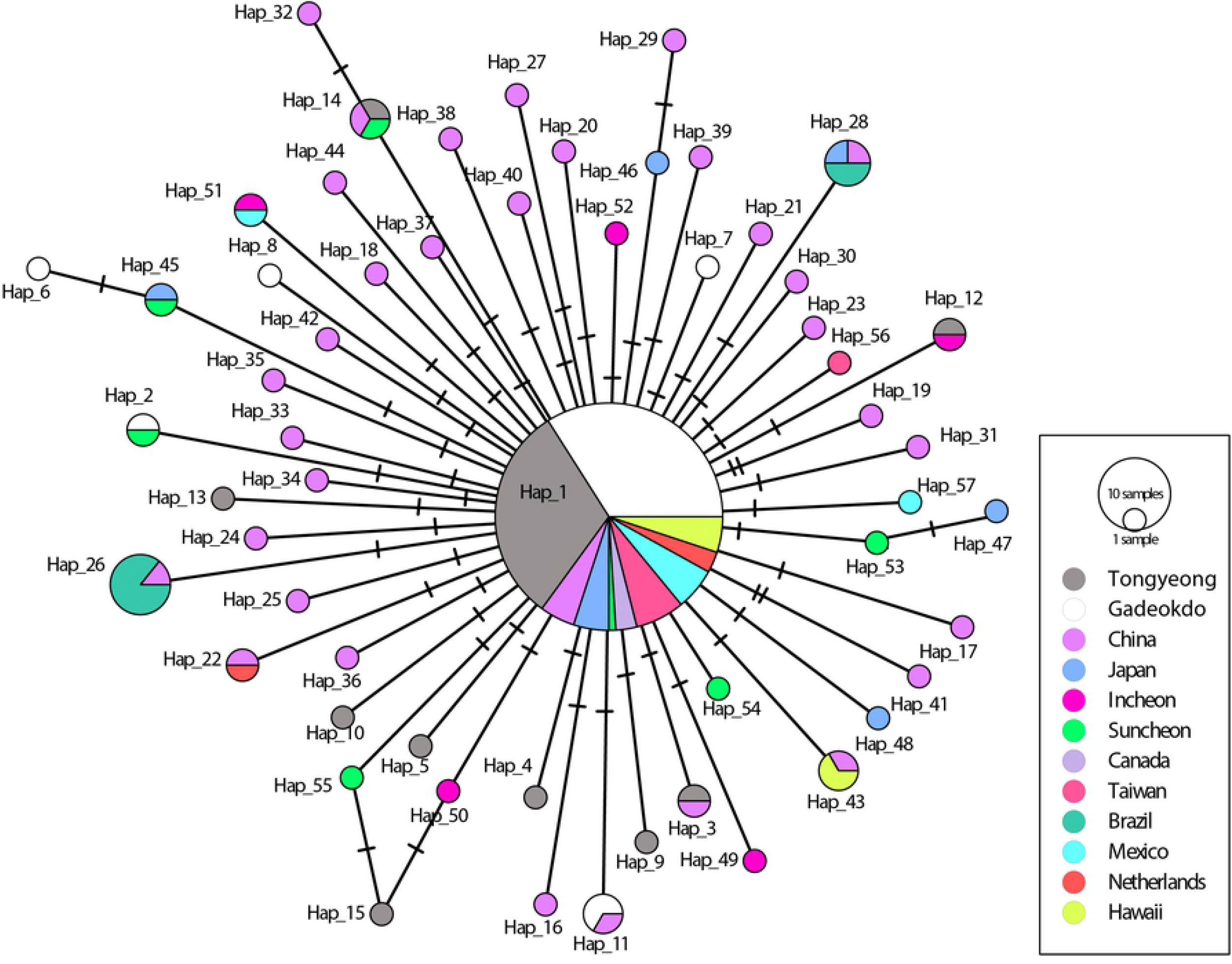
Haplotype network of mtCOI of 12 *M. gigas* populations, including TY and GD. Each of the different colors represents a geographic sampling site. Each circle corresponds to a unique haplotype sized in proportion to its frequency. The hash marks on the lines indicate the number of substitutions that separate haplotypes.

## Discussion

The loss of genetic diversity reduces fitness and adaptability of a species and consequently the size of its population, increasing the risk of extinction of the species. Rapid decline of *M. gigas* populations in recent years necessitates a science-based fisheries management in order to increase the gene pool and sustainability of *M. gigas* populations. To better understand the genetic diversity of *M. gigas*, two neighboring aquaculture farms, Tongyeong and Gadeokdo, located on the southern coast of Korea were investigated in the present study. Both *M. gigas* populations (TY and GD, hereafter) were shown to possess low levels of genetic diversity, which is considered typical to the extant Pacific oyster *M. gigas* populations [21, 23]. Our data also suggest that fine-scale spatial and temporal genetic characteristics exist in these populations and a full understanding of such system requires further study in the historical, biological, and environmental contexts.

### Genetic features of *M. gigas* populations

A total of 21 haplotypes were identified in TY and GD populations, based on SNPs within the mtCOI gene. The subsequent genealogical analysis showed that these haplotypes form a characteristic shallow network consisting of the major haplotype (i.e., Hap_1) at the center and 20 minor satellite haplotypes diverging from HAP_1 (Fig. 3). Genetic differences between the two populations were shown to lie in their distinctive satellite haplotypes; TY had 12 satellite haplotypes while GD had 8, but none of the satellite haplotypes were shared between the two. Each satellite haplotype differed from HAP_1 by 1-3 mtCOI SNPs and mostly formed a singleton, suggesting these satellite haplotypes have emerged recently. Previously, Li et al. reported the similar “start-like” network based on 228 mtCOI sequences of wild *M. gigas* collected from the coasts of China, Korea, and Japan [21]. Furthermore, HAP_1 accounted for more than 70% of both wild and cultured populations, suggesting the population genetic structure of *M. gigas* is conserved between wild and hatchery populations. The sharing of HAP_1 at equally high frequencies among the geographically diverse populations indicates that HAP_1 likely represents the ancestral spat oyster from Japan that has become prevalent worldwide since the early 20^th^ century [1]. Weak genetic differentiation (FST = 0.00188) between TY and GD is, thus, more likely due to the history of monophyly and large-scale distribution associated with the current *M. gigas* industry, rather than distance-dependent genetic homogenization between the two populations.

### Spatial and temporal relatedness among *M. gigas* populations

*M. gigas* populations were further characterized by analyzing a haplotype network of 117 mtCO1 sequences from 12 different geographic origins (Fig 4). This extended network consisted of the predominant HAP_1 and 56 satellite haplotypes, a majority of which being a singleton. Geographic clustering among the 4 Korean locations (TY, GD, Incheon, and Suncheon) and among neighboring nations (China, Korea, and Japan) was not observed, as expected given the large number of singletons in the network. Only 3 satellite haplotypes (i.e., HAP_3, HAP_11, and HAP_14) were shared between analyzed Chines and Korean *M. gigas* populations. This data further supports the notion that geographic proximity has minimal influence on the genetic make-up of *M. gigas* populations. Therefore, it is plausible that gene flow between TY and GD populations is not significant whereas genetic drift is the main mechanism for genetic diversification of each population. This is surprising given the ocean ecology of these two regions is believed to be commonly affected by the Kuroshio current [34]. The finding that TY had slightly higher variation than GD suggests that fine-scale environmental factors might play a role in the equilibrium between gene flow and genetic drift, which needs further study [35].

As the samples from TY and GD were collected over the four seasons, we examined temporal correlation between season and haplotype diversity. Distinct haplotypes constituted each season, and frequent replacements of haplotypes were detected over the 4 seasonal periods except for Hap_1 (Table 4). This data suggests that genetic variation was not passed down from one season to the next (i.e., the absence of self-recruiting). This result may be due to the insufficient sampling size to detect recruited haplotypes, but maybe explains how a novel genetic variant can get lost over time within a population. In addition, the spring cohort had the lowest level of genetic variation (one satellite haplotype) in both populations, indicating that the rate of genetic variation fluctuates over time and may be influenced by seasonal selection related to reproductive activity, food availability, water temperature, etc [36, 37].

Altogether, our findings provide insight into genetic diversity and spatial and temporal dynamics of *M. gigas* populations on the coastal aquafarms in Korea. Further research in this topic should help improve genetic monitoring and sustainable management of *M. gigas* populations.

## Conclusions

In the present study, the Pacific oyster *M. gigas* of two neighboring aquafarms in Korea were investigated for the genetic diversity and population structure inferred by mtDNA CO1 sequences. Consistent to the monophyletic nature of this species, both *M. gigas* populations were shown to have a similar genetic structure and low genetic diversity. Comparative haplotype analyses provided data supporting new genetic features of the populations that include (1) weak genotype-locality relationship, (2) low levels of gene flow between populations, and (3) seasonal fluctuation of genetic variation within a population. These results indicate genetic dynamics of *M. gigas* populations are complex and warrant further study in biological and ecological contexts. Information from such study will be the groundwork to set strategies for reducing genetic erosion and developing more sustainable management and breeding program for *M. gigas* populations.

